# A UHM – ULM interface contributes to U2AF2 and SF3B1 association for pre-mRNA splicing

**DOI:** 10.1101/2022.01.09.475535

**Authors:** Justin Galardi, Victoria N. Bela, Nazish Jeffery, Xueyang He, Eliezra Glasser, Sarah Loerch, Jermaine L. Jenkins, Mary J. Pulvino, Paul L. Boutz, Clara L. Kielkopf

## Abstract

In the early stages of spliceosome assembly, the 3’ splice site is recognized by sequential complexes of U2AF2 with SF1 followed by the SF3B1 subunit of the U2 small nuclear ribonucleoprotein particle. The U2AF2 – SF1 interface comprises a U2AF homology motif (UHM) of U2AF2 and a well-characterized U2AF ligand motif (ULM)/coiled coil region of SF1. However, the structure of the U2AF2 – SF3B1 interface and its importance for pre-mRNA splicing is unknown. To address this knowledge gap, we determined the crystal structure of the U2AF2 UHM bound to a SF3B1 ULM site at 1.8 Å resolution. The trajectory of the SF3B1 ULM across the U2AF2 UHM surface differed from prior UHM/ULM structures. This distinctive structure is expected to modulate the orientations of the fulllength proteins. Using isothermal titration calorimetry, we established similar binding affinities of a minimal U2AF2 UHM – SF3B1 ULM complex and a nearly full-length U2AF2 protein binding the N-terminal SF3B1 region, with or without an auxiliary SF3B6 subunit. We showed that key residues at the U2AF2 UHM – SF3B1 ULM interface are required for high affinity association and co-immunoprecipitation of the splicing factors. Moreover, disrupting the U2AF2 – SF3B1 interface altered splicing of representative human transcripts. Further analysis of these transcripts and genome-wide data sets indicated that the subset of splice sites co-regulated by U2AF2 and SF3B1 are largely distinct from those co-regulated by U2AF2 and SF1. Altogether, these findings support distinct structural and functional roles for the sequential SF1 and SF3B1 complexes with U2AF2 during the pre-mRNA splicing process.

## Introduction

The spliceosome assembles on consensus splice site signals of the pre-mRNA in a series of ATP-dependent conformational transitions (reviewed in (1)). In the initial ATP-independent E-complex, the essential pre-mRNA splicing factor U2AF2 recognizes a polypyrimidine (Py) tract preceding the 3’ splice site (2–4) as a heterodimer with a U2AF 1 small subunit, which contacts an AG at the splice site junction (5). First, U2AF2 forms a ternary complex with SF1 (6,7), which in turn recognizes the branch point consensus sequence (BPS) (8). In the subsequent A-complex, U2AF2 recruits the U2 small nuclear ribonucleoprotein particle (snRNP) of the spliceosome to the 3’ splice site. At this stage, the SF3B1 spliceosome subunit of the U2 snRNP replaces SF1 in the U2AF2 complex (9,10). Following several ATP-dependent conformational changes among the core snRNP particles and dissociation of U2AF2 (10,11), the spliceosome ultimately achieves the activated B^ACT^-complex. A final conformational change to the B*-complex allows the first catalytic reaction of pre-mRNA splicing to generate the branched intron lariat.

This parts list of core spliceosome assemblies has been illuminated by recent cryo-electron microscopy (cryoEM) structures of B, B^ACT^, C, C* and intron-lariat spliceosomes (reviewed in (12)). Nevertheless, cryoEM approaches have yet to resolve the early stages of 3’ splice site recognition, which is challenging due to a fleeting “dance” of transitions among low molecular mass subunits. A cryoEM structure of a 5’ splice site in the E-like yeast spliceosome assembly revealed weak density that could not be reliably modeled as the U2AF2 and SF1 homologues (Mud2 and BBP) (13). Although SF3B1-containing structures of spliceosomes are available (14–19), the U2AF2 – SF3B 1 complex has not been resolved. As such, the field’s structural understanding of U2AF2 and its partners remains limited to piecewise structures of the interacting domains.

A C-terminal “U2AF Homology Motif” (UHM) domain of U2AF2 binds a well-characterized “U2AF Ligand Motif” (ULM) adjoining a coiled-coil region of SF1 (20,21). Such UHM family members are marked by an RNA recognition motif (RRM)-like fold with specialized features for recognizing ULM proteins, as opposed to RNA (reviewed in (22)). Several UHM proteins are known to associate with five ULMs in an intrinsically unstructured, N-terminal region of SF3B1, which additionally contains two ULM-like motifs without known UHM partners (**Fig. 1*A***). *In vitro* assays with purified proteins have shown that the U2AF2 UHM binds the SF3B1 ULMs (23–25). The highest affinity U2AF2 UHM binding site is the fifth SF3B1 ULM (ULM5) (23), which is adjacent a binding site for an SF3B6 subunit. However, the structure of the U2AF2 – SF3B1 complex and its relevance for pre-mRNA splicing is unknown.

**Figure 1.**
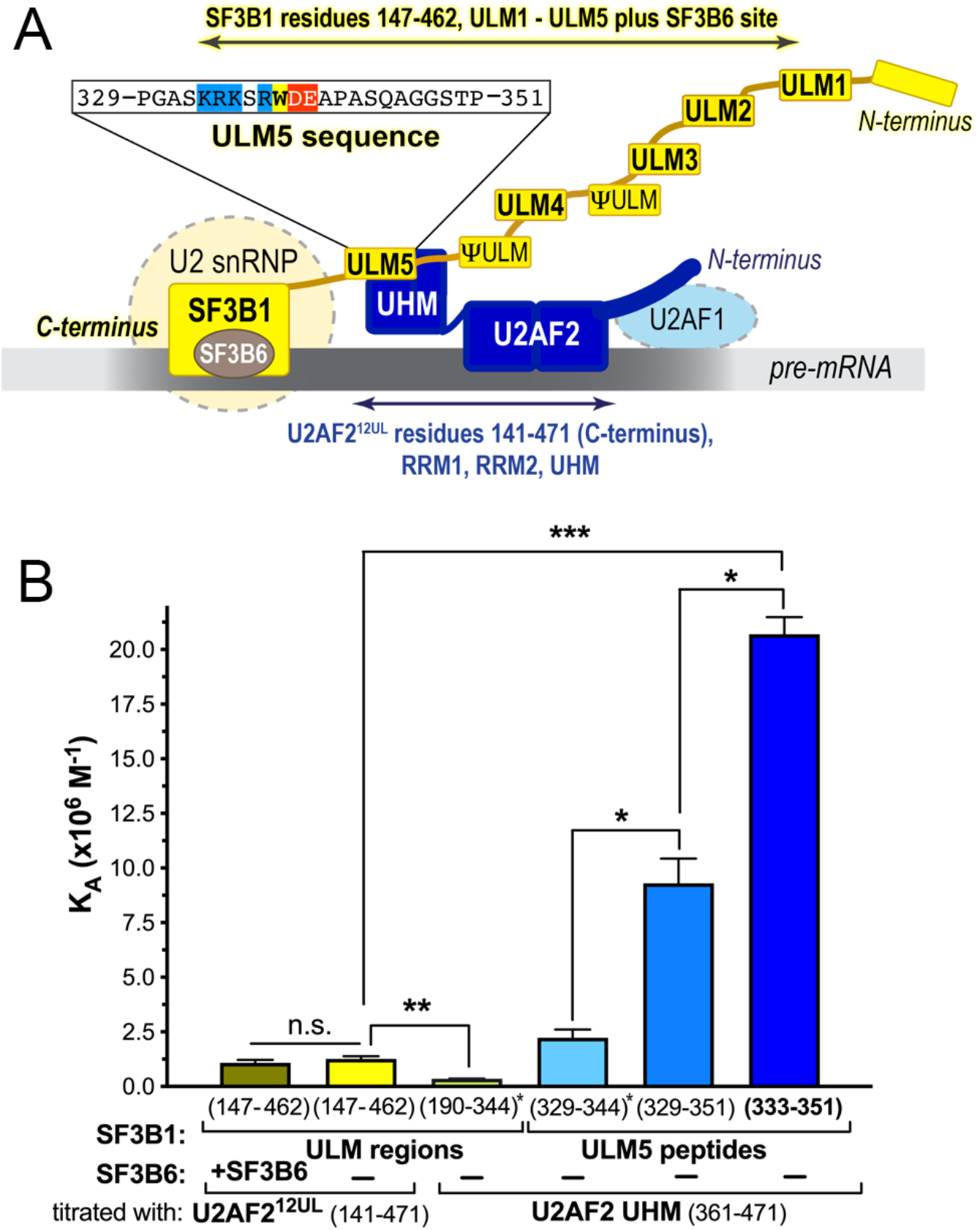
SF3B1 ULM5 (residues 333-351) is a relevant, high-affinity ligand for the U2AF2 UHM. *A,* Schematic diagram of the SF3B1 (yellow) – U2AF2 (navy) complex at the 3’ splice site (gray). The sequence of the ULM5 region is expanded above, with the ULM consensus marked in color for basic residues (blue), acidic residues (red), or tryptophan (yellow). Boundaries of regions used for ITC are indicated by double-headed arrows. *B,* Bar graph of the average apparent binding affinities (K_A_) and standard deviations of three isothermal titration calorimetry experiments. The residue ranges of the SF3B1 fragment in the sample cell are given on the x-axis and the U2AF2 residues are listed below. SF3B6 (also called p14a) is included in the first experiment of the graph (taupe). Values reproduced from Thickman *et al* (2006) *J. Mol. Biol.* v356:664 are marked by asterisks. The thermodynamic values and representative isotherms are given Table S1 and Fig. S1. Significance was assessed by unpaired t-tests with Welch’s correction implemented in GraphPad Prism, here and for all figures: n.s., not significant; *, p<0.05; **, p<0.005; ***, p<0.0005.

Here, we determine the U2AF2 UHM – SF3B1 ULM5 crystal structure. The ULM5 ligand conformation diverges from prior UHM – ULM complexes, highlighting the importance of determining new structures in the UHM – ULM family. We find that the SF3B6 subunit has no detectable influence on the U2AF2 – SF3B1 binding affinity. We demonstrate the functional importance of the U2AF2 UHM – SF3B1 ULM5 interface for association of these splicing factors and provide evidence for its functional contributions to splicing of pre-mRNA transcripts.

## Results

### SF3B1 ULM5 is a high affinity binding site for U2AF2 that is independent of SF3B6

We used isothermal titration calorimetry (ITC) to test whether extending the SF3B1 region to N-terminal residues preceding ULM1, including the C-terminal SF3B6 binding site, and adding the SF3B6 subunit, would alter the U2AF2 binding affinity (**Fig. S1**). Additionally, we extended the boundaries of the U2AF2 construct (U2AF2^12UL^) to the nearly full-length protein except for an unstructured N-terminal region that binds U2AF1 (**Fig. 1*A***). The affinities of the lengthened proteins increased slightly (by three-fold) compared to the U2AF2 UHM binding a prior, shorter SF3B1 construct (**Fig. 1*B***). The presence of SF3B6 had no significant effect on the binding affinity of SF3B1 for U2AF2^12UL^, supporting a prior result that the binding sites for these subunits do not overlap (24). More notably, the apparent stoichiometry decreased to two U2AF2^12UL^ per SF3B1 ULM region, rather than the three-to-one stoichiometry observed previously for the U2AF UHMs (**Table S1**). The smaller size of the U2AF2 UHM than U2AF2^12UL^ is likely to leave room for a third U2AF2 molecule to associate with the SF3B1 ULMs. We note that the ability of excess U2AF2^12UL^ to concurrently bind more than one of the SF3B1 ULMs does not necessarily reflect the stoichiometry of full-length U2AF2 in the context of the assembling spliceosome, which is thought to contain a single U2AF2 and U2 snRNP per splice site (26).

We next compared the U2AF2 UHM affinities for binding SF3B1 ULM5 peptides with different boundaries. The SF3B1 ULM5 site has the highest apparent affinity among SF3B1 ULM variants binding a tryptophan-less U2AF2 UHM in intrinsic tryptophan fluorescence experiments (23). By ITC, the U2AF2 UHM binds a ULM5 peptide (residues 329-344) with slightly higher affinity than its average apparent affinity for all sites in an SF3B1 ULM region (residues 190-344) (23). Extending the ULM5 C-terminus to include a TP-motif, which is a phosphorylation site in human cells (27,28), increased its U2AF2 UHM binding affinity by fourfold (**Fig. 1**, **Table S1**). On the other hand, truncating the ULM N-terminus to remove residues preceding an R/K-rich motif slightly increased the U2AF2 UHM affinity by two-fold. These results defined a high affinity ULM5 region (residues 333-351) for co-crystallization with the U2AF2 UHM.

### The SF3B1 ULM5 binds to the U2AF2 UHM in an atypical trajectory

To view the U2AF2 UHM – SF3B1 ULM5 interactions, we determined the crystal structure at 1.80 Å resolution (**Table S2**). The crystallographic asymmetric unit contained two similar copies of the U2AF2 UHM – SF3B1 ULM5 complex (RMSD 1.45 Å for 113 matching Ca atoms) (**Fig. S2*A***). The electron density maps revealed twelve or nine ordered residues for the non-crystallographic symmetry (NCS)-related copies of UHM-bound SF3B1 ULM5 (**Fig. S2*B***). The additional ordered residues append a C-terminal α-helical turn (residues 344-346) that makes crystallographic and NCS-related contacts with neighboring UHMs.

The central tryptophan (W338) of each ULM5 copy is inserted between the UHM a-helices (**Fig. 2*A***). The W338 side chain is sandwiched in a T-type interaction between F454 and an R452 – E405 salt bridge. The R337 side chain further anchors the SF3B1 ULM by a distinct salt bridge with the U2AF2 UHM E397. Additionally, disordered basic residues at the N-terminus of the ULM5 are aligned for electrostatic attraction with the acidic UHM α-helix. These prominent ULM5-interacting residues of U2AF2 UHM are located in an RXF motif (F454, R452) and acidic a-helix (E405, E397) that are characteristic of the UHM family (22), as illustrated for RBM39 (also called CAPERa) bound to SF3B1 ULM5 (29) (**Fig. 2*B***).

**Figure 2.**
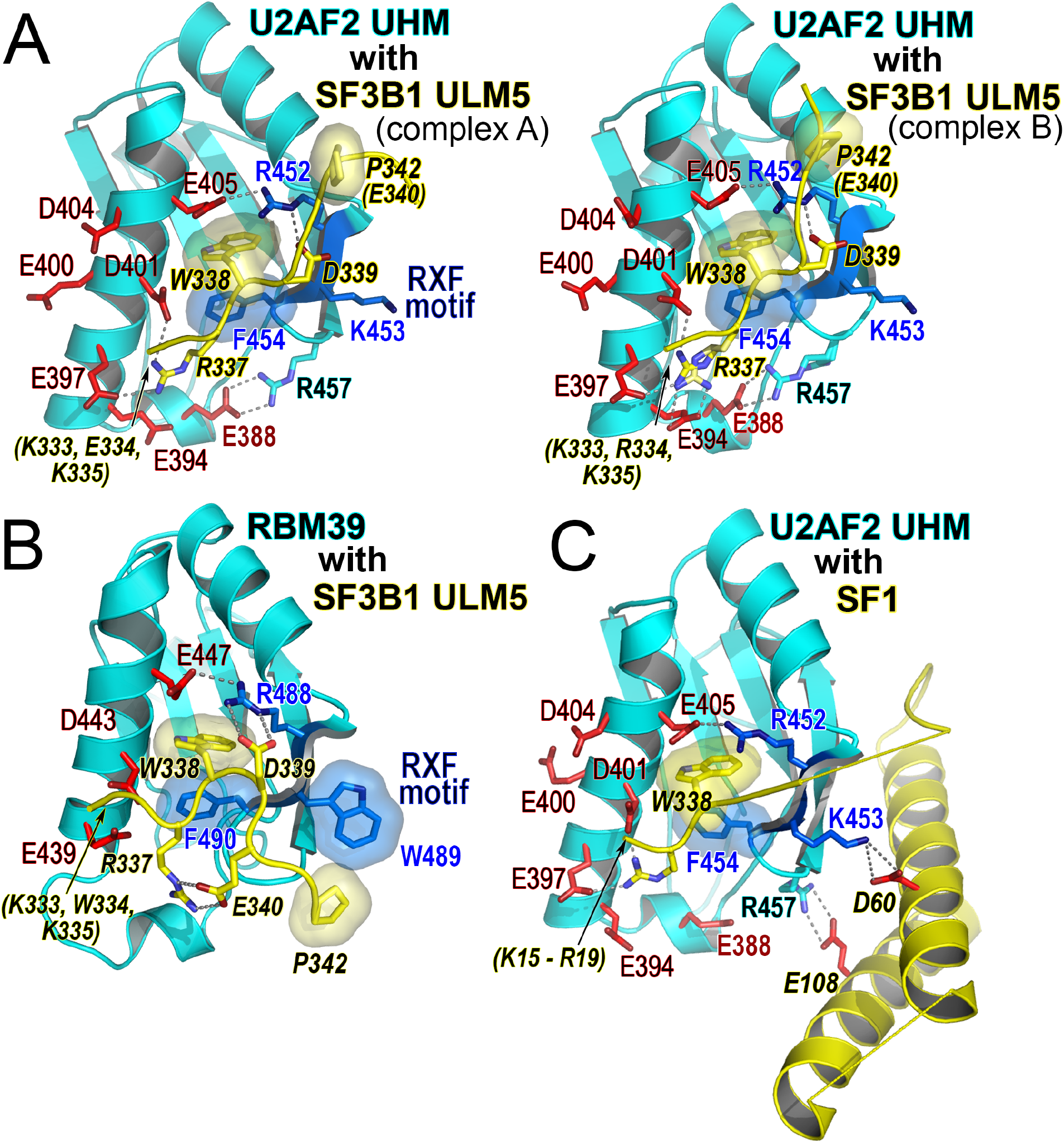
Structure of the U2AF2 UHM bound to SF3B1 ULM5 and comparison with related complexes. *A,* Views of the two U2AF2 UHM – SF3B1 ULM 5 complexes in the crystallographic asymmetric unit. The R337 and R457 side chains of complex B have two alternative conformations*. B,* RBM39 UHM – SF3B1 ULM5 (PDB ID 4OZ1) or *C,* U2AF2 UHM – SF1 (PDB ID 4FXW) complexes viewed in a similar orientation. The UHM is colored cyan; the ULM ligand is yellow with italicized residue labels; key acidic UHM residues are red; residues in the UHM RXF motif are blue; and disordered residues are labeled in parentheses.

Both SF3B1 ULM5 copies follow a linear trajectory that runs nearly parallel to the acidic α-helix of the U2AF2 UHM (**Fig. 2*A***). The proline of the ULM5 TP-motif packs against the UHM surface in a depression above the R452 – E405 salt bridge and preceding the RXF loop. This relatively straight conformation of the ULM backbone differs from other UHM – SF3B1 ULM complexes characterized to date (e.g., RMSD 2.9 Å between eight matching Ca of the RBM39-bound SF3B1 ULM5 in **Fig. 2*B***). More typically, U-shaped SF3B 1 ULMs pack the TP motif against an exposed aromatic side chain from the central “X” position of the UHM RXF motif (e.g., W489 of RBM39 in **Fig. 2*B***). The U2AF2 UHM lacks this aromatic residue, which instead is replaced by K453.

In the SF1 complex, this K453 residue of the U2AF2 RXF motif mediates a salt bridge with the SF1 ULM-coiled coil region (20,21). The U2AF2 R457 side chain forms a second salt bridge with the SF1 coiled coil and also encloses F454 below the central ULM tryptophan (**Fig. 2*C***). Although the angle of the SF1 ULM polypeptide relative to the UHM a-helix is greater than for SF3B1 ULM, the backbone conformations are similar (RMSD 0.5 Å for five matching Ca of SF1 ULM and SF3B1 ULM5). Like the U2AF2-bound SF3B1 ULM5 and distinct from other UHM – ULM complexes, the C-terminal residues of the SF1 ULM are located closer to R452 than to the central position of the RXF motif. Nevertheless, the SF1 coiled coil makes little contribution to its binding affinity for U2AF2 (23,30) and instead may influence the specific orientation of the SF1 and U2AF2 domains. Indeed, the isolated SF1 ULM has high affinity for the U2AF2 UHM (24 nM), possibly due to extension of the basic tail (six residues for the SF1 ULM compared to four residues for SF3B1 ULM5). The comparable U2AF2 binding affinities of SF1 and SF3B1 ULM5 invoke roles for other factors to prompt exchange of SF1 for SF3B1 during spliceosome assembly.

### Interface mutations reduce U2AF2 UHM – SF3B1 ULM5 binding affinity

We probed key residues at the U2AF2 UHM – SF3B1 ULM interface by ITC of structure-guided mutant proteins (**Fig. 3**, **Fig. S1**, **Table S1**). The amino acid substitutions are unlikely to perturb the overall protein folds since we targeted residues at the UHM surface and the ULM is unstructured (23). First, we tested the relevance of UHM contacts with the C-terminal residues of the ULM. Since SF3B1 T341 has an intramolecular contact that positions M346 in complex A, we investigated a T341A/M346A double mutant. We also tested a glycine substitution for P342 that is expected to confer a flexible backbone without a side chain, in contrast with the wild-type proline. The U2AF2 UHM binding affinities of the T341A/M346A and P342G SF3B1 ULM5 mutants were similar to the wild-type (WT) counterpart, consistent with the variable positions of these residues in the two crystallographically-independent copies of the complex.

**Figure 3.**
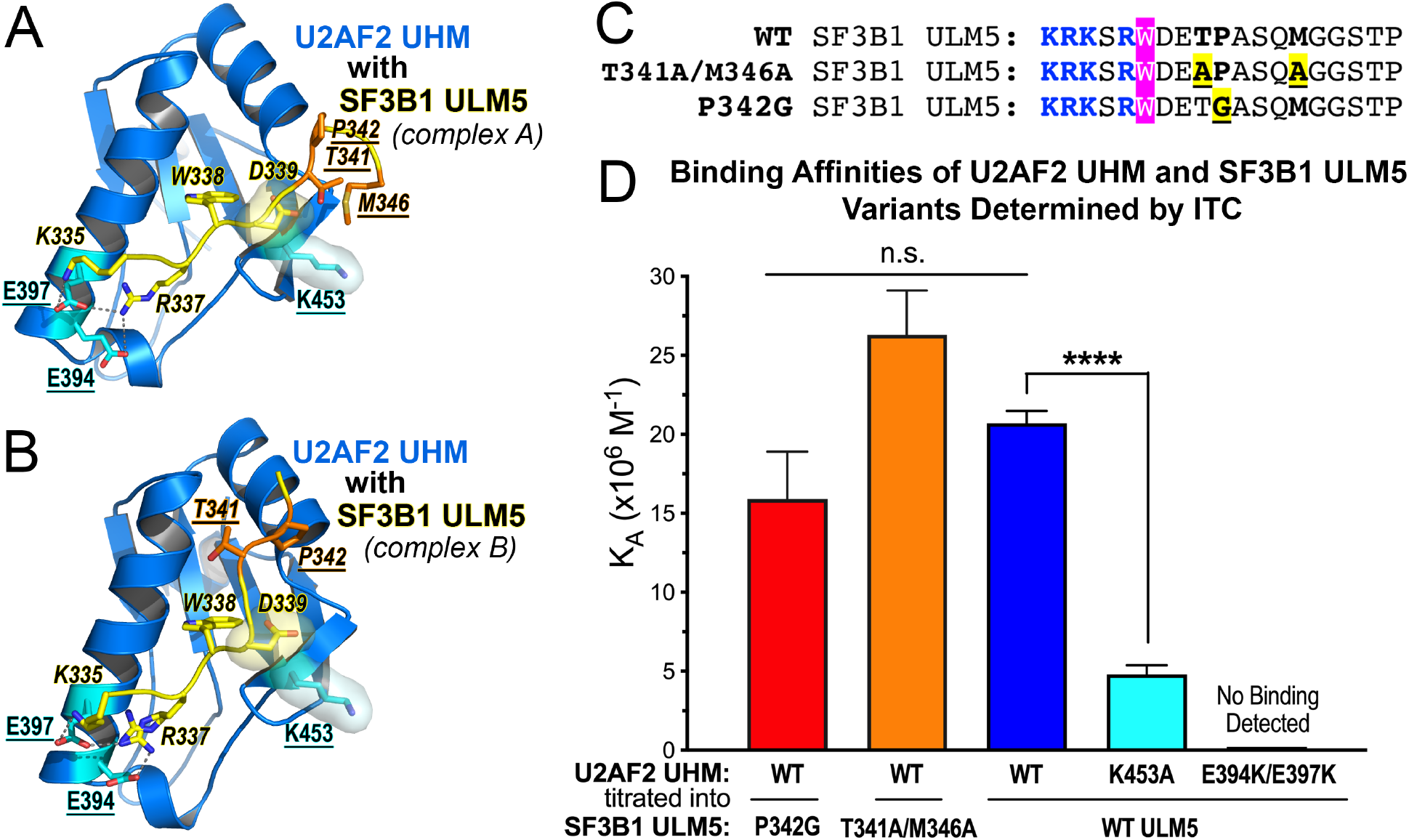
Contribution of interface residues to the U2AF2 UHM – SF3B1 ULM5 binding affinity. *A, B*, Interactions mediated in the two copies of the U2AF2 UHM (marine) – SF3B1 ULM5 (yellow, italicized labels) complex tested by structure-guided mutagenesis (underlined labels; U2AF2: cyan; SF3B1: orange). *C,* Sequences of the SF3B1 ULM5 peptides (residues 333-351) showing the structure-guided mutants (yellow) tested by ITC. The central tryptophan is magenta and the basic residues are blue. *C,* Bar graph of the average apparent binding affinities (K_A_) and standard deviations of three isothermal titration calorimetry experiments. The thermodynamic values and representative isotherms are given Table S1 and Fig. S1. n.s., not significant; ****, p<0.0001.

Next, we tested a potential role for the U2AF2 K453 residue at the central position of the UHM RXF motif. This U2AF2 lysine differs from a typically aromatic residue at this position of other UHMs, which appears to influence the bound ULM conformation (**Fig. 2**). The distinct trajectory of the U2AF2-bound ULM5 packs SF3B1 D339 against the hydrophobic portion of the U2AF2 K453 side chain (**Fig. 3*A-B***). Accordingly, replacing K453 with alanine reduced the binding affinity of the U2AF2 UHM – SF3B1 ULM5 complex by four-fold (**Fig. 3*D*, Table S1**). This small, but statistically significant, change is consistent with loss of the observed D339 – K453 contact and/or disruption of a weak ionic interaction between the side chains.

Lastly, we examined interactions between the canonical acidic a-helix of the UHM and basic residues of the N-terminal ULM tail (**Fig. 3*A-B***). Substituting lysines for the U2AF2 E394/E397 residues abolished detectable binding to the SF3B1 ULM5 (**Fig. 3*D***, **Table S1**). This result, coupled with our prior observations that the central ULM tryptophan is required for detectable association of the purified U2AF2 UHM and SF3B1 proteins (23,25), equipped us with structure-guided mutants to investigate the functional importance of this interface.

### The UHM-ULM interface contributes to U2AF2 – SF3B1 association in human cells

We investigated whether association of the fulllength U2AF2 and SF3B1 proteins in coimmunoprecipitations from human cells (HEK 293T) relied on the UHM – ULM interface (**Fig. 4**). The N-terminally tagged constructs for U2AF2 (HA-tag, ^HA^U2AF2) and SF3B1 (FLAG-tag, ^FLAG^SF3B1) were transiently co-expressed in HEK 293T cells. The ^HA^U2AF2-associated protein complexes were immunoprecipitated using anti-HA agarose beads. We found that wild-type ^FLAG^SF3B1 efficiently associated with ^HA^U2AF2 (**Fig. 4*B***). The ^HA^U2AF2 E394K/E397K mutation abolished detectable co-immunoprecipitation of ^FLAG^SF3B1, consistent with the ability of this mutation to disrupt U2AF2 UHM binding to the SF3B1 ULM-containing region in ITC experiments.

**Figure 4.**
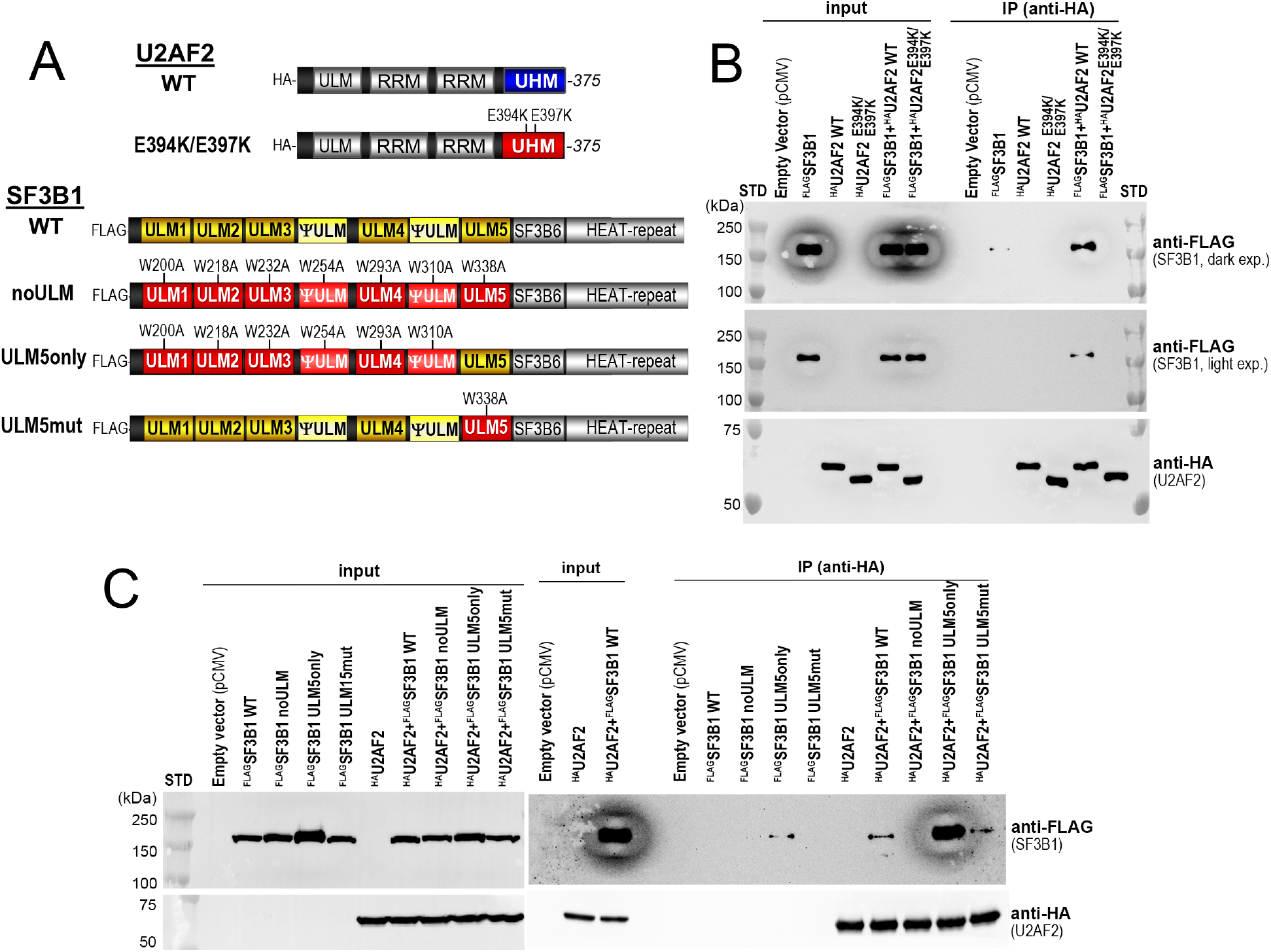
The U2AF2 UHM contributes to association with SF3B1 in human cell extracts. Coimmunoprecipitation showing the amounts of FLAG-tagged SF3B1 (^FLAG^SF3B1, 148 kDa) retained by HA-tagged U2AF2 (^HA^U2AF2, 55 kDa) variants co-expressed in HEK 293T cells. Associations between the wild-type (WT) and structure-guided mutant proteins diagrammed in *A* were compared, including *B,* a E394K/E397K U2AF2 UHM-mutant, or *C,* SF3B1 ULM mutants. “Input” controls correspond to 0.5% and 1% of the total protein amounts used for the co-immunoprecipitations shown in *B* and *C*. STD, molecular size markers.

We then examined the effects of mutating the SF3B1 ULM tryptophans to alanine, which is known to prevent detectable U2AF2 – SF3B1 binding in ITC experiments (23). The ^FLAG^SF3B1 mutations either affected all ULMs and pseudo– ULMs (“noULM”), left only ULM5 intact (“ULM5only”), or disrupted only ULM5 (“ULM5mut”) (**Fig. 4*A***). As expected considering ITC results (23), the ^FLAG^SF3B1 noULM variant no longer detectably co-immunoprecipitated with ^HA^U2AF2 (**Fig. 4*C***). Conversely, ^HA^U2AF2 association with ^FLAG^SF3B1 ULM5only appeared to significantly increase in the absence of the other intact ULMs/pseudo-ULMs. Since the ULM5only variant and wild-type SF3B1 ULM regions have similar apparent affinity for the U2AF2 UHM in ITC experiments with recombinant proteins, this enhanced interaction is likely due to disruption of additional regulatory factors present in cells. Following mutation of only ULM5, the ^FLAG^SF3B1 noULM5 protein continued to co-immunoprecipitate with similar amounts of ^HA^U2AF2 as wild-type ^FLAG^SF3B1, consistent with the ability of other SF3B1 ULMs to bind the U2AF2 UHM (23).

### The U2AF2 UHM – SF3B1 ULM interface contributes to splicing of representative transcripts

The UHM – ULM-dependent association of U2AF2 with SF3B1 suggested that this interface could contribute to the pre-mRNA splicing functions of these proteins. To test this hypothesis, we first made use of a well-characterized, U2AF2-sensitive minigene comprising alternative 3 splice sites *(py* and *PY)* (31,32) (**Fig. 5*A***). As noted previously in our cell culture conditions (32), HEK 293T cells stably expressing the *pyPY* minigene produced mostly unspliced *pyPY* transcript by reverse-transcription (RT)-PCR (**Fig. 5*B***). Overexpression of wild-type U2AF2 increased splicing of the *py* site substantially and the *PY* splice site moderately by RT-PCR and quantitative real-time (q)RT-PCR (**Fig. 5*B-C***). By contrast, most of the *pyPY* transcript remained unspliced following overexpression of the E394K/E397K mutant U2AF2. These results confirmed that the U2AF2 UHM contributes to 3’ splice site selection for the *pyPY* prototype.

**Figure 5.**
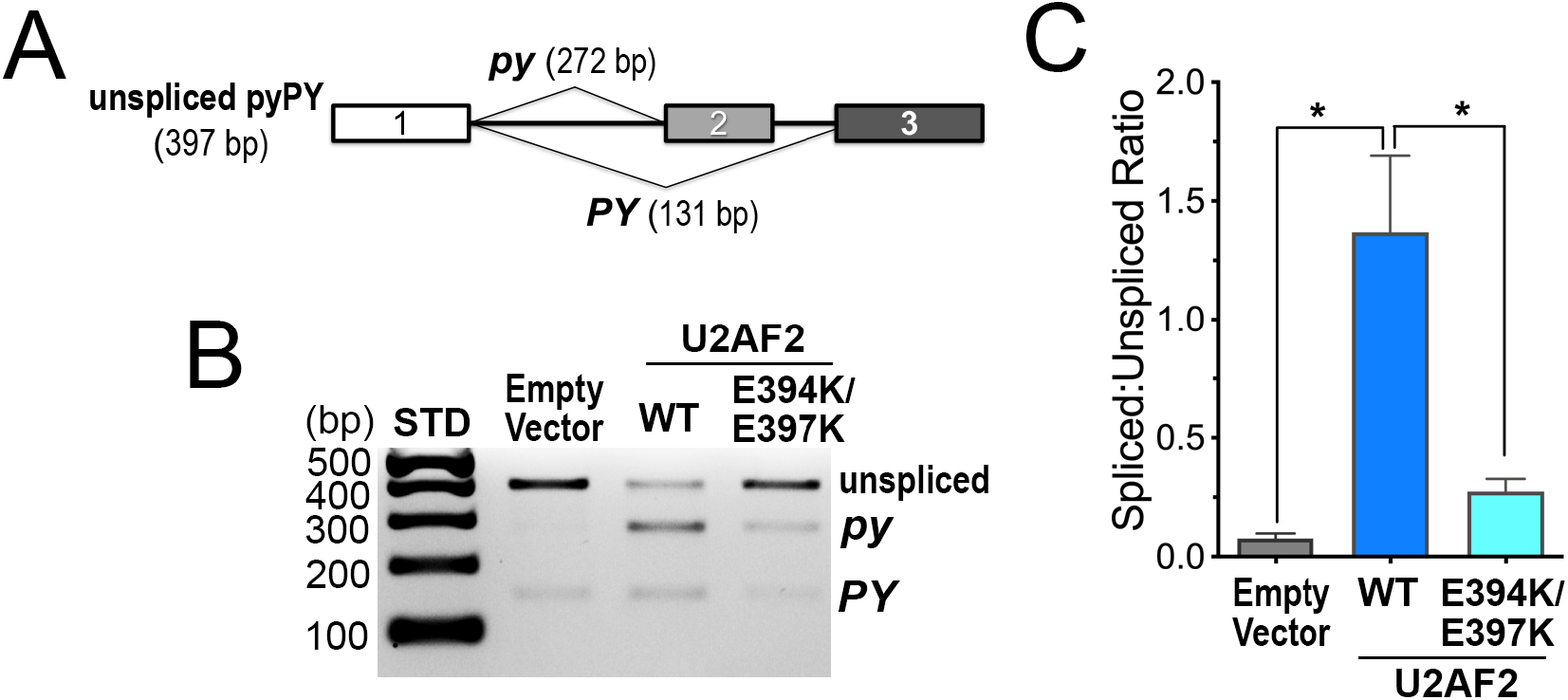
The U2AF2 UHM influences splicing of a prototypical minigene substrate. *A,* Schematic diagram of the *pyPY* transcript. Alternatively spliced sites generate two differently-sized products *(py,* 272 bp and *PY*, 131 bp). The unspliced*pyPY* RT-PCR product is 397 bp. *B,* Representative RT-PCR of total RNA isolated from HEK293T cells stably expressing the *pyPY* minigene and transfected with either wild-type (WT) or a double mutant U2AF2 (E394K/E397K). The RT-PCR products were resolved by agarose gel electrophoresis with ethidium bromide staining. *C,* Quantification o f*pyPY* RT-PCR products. The band intensities from three independent replicates were background-corrected and averaged. The average ratios of the total spliced product *(py* + *PY)* to the unspliced *pyPY* and standard deviations are plotted for each sample. Immunoblots are shown in Fig. S3. STD, 100 bp ladder molecular size markers. *, p<0.05.

In the next step, we asked whether SF3B1 and the SF3B1 ULMs contribute to alternative splicing of endogenous, U2AF2-responsive transcripts. We and others have shown that skipping of *THYN1, SAT1, INTS13,* and *RNF10* exons is sensitive to reduced U2AF2 levels (32–34). Although we were unable to achieve robust rescue to test the effects of structure-guided U2AF2 variants (32), these precedents offer a means to compare the potential contributions by the SF3B1 ULMs to U2AF-sensitive splicing events.

In support of inter-related U2AF2 and SF3B1 functions, siRNA-mediated reductions in SF3B1 levels increased exon-skipped splicing of these transcripts in a similar manner as U2AF2 knockdown (**Fig. 6, Fig. 7**).

**Figure 6.**
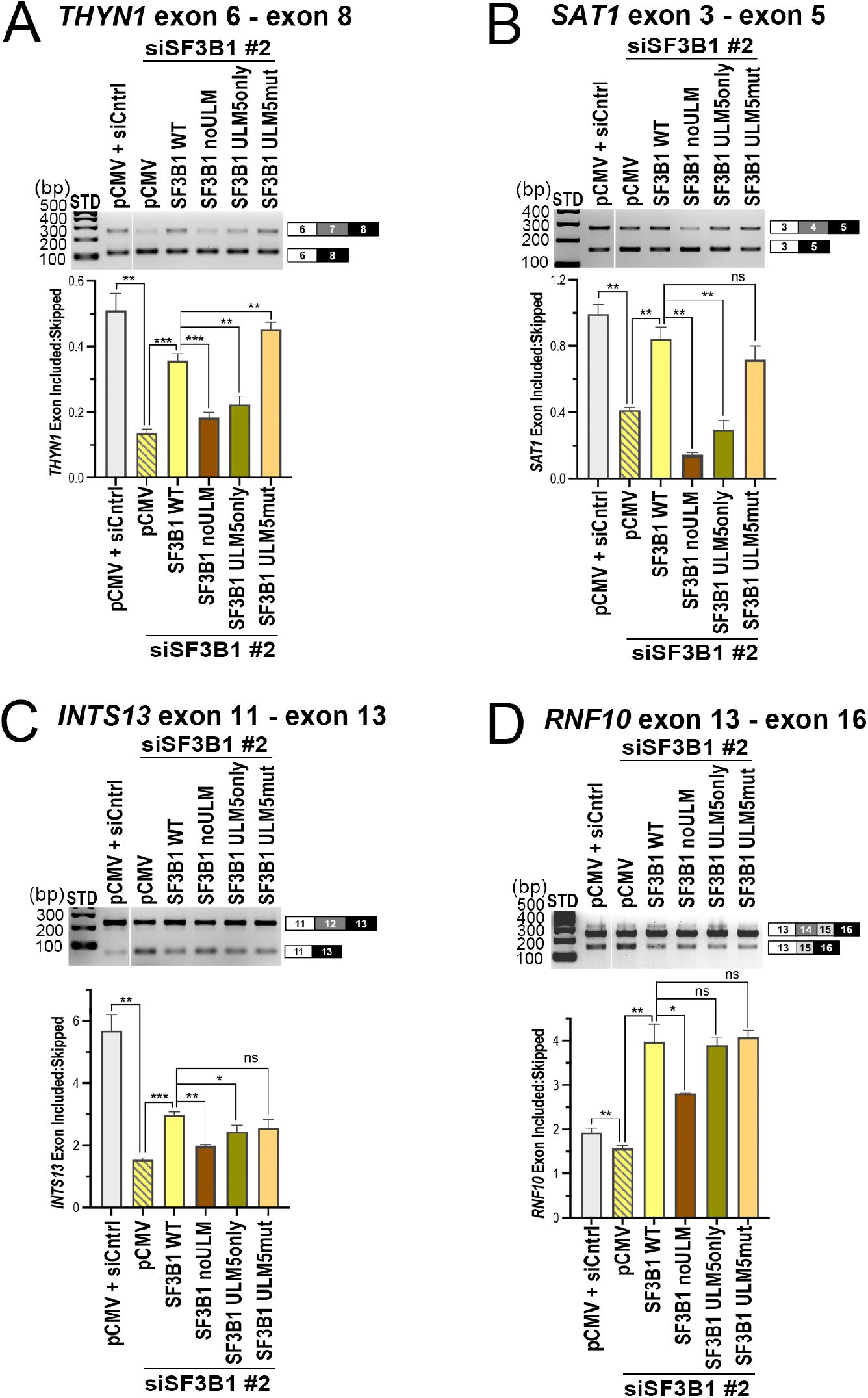
The SF3B1 ULMs contribute to accurate splicing of endogenous, U2AF2-responsive gene transcripts. The abilities of indicated SF3B1 ULM variants to rescue splicing of representative U2AF2-regulated transcripts, including *A, THYN1; B, SAT1; C, INTS13;* and *D, RNF10,* were compared by RT-PCR. The ethidium bromide-stained agarose gels are shown above. The ratios of exon-included:exon-skipped band intensities for background-corrected and averaged replicates were plotted below. Primer sequences are listed in Table S3, and immunoblots are shown in Fig. S4. STD, 100 bp ladder size standards. n.s., not significant; *, p<0.05; **, p<0.005; ***, p<0.0005.

**Figure 7.**
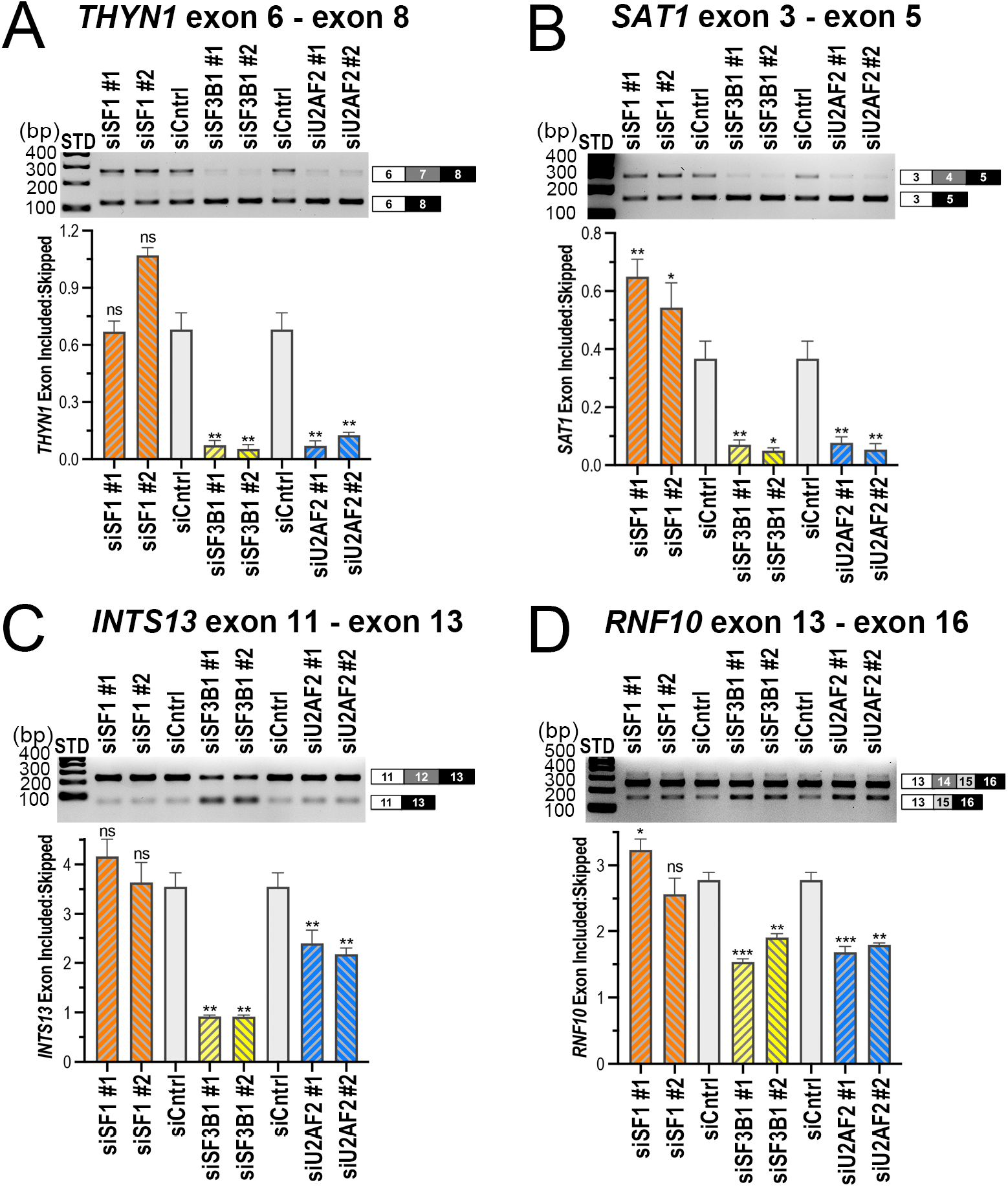
Comparison of SF3B1 and SF1 modulation of representative U2AF2-responsive gene transcripts. The effects of siRNA-mediated reductions in SF1, SF3B1, or U2AF2 levels on splicing of representative U2AF2-regulated sites, including *A, THYN1; B, SAT1; C, INTS13;* and *D, RNF10,* were compared by RT-PCR as described for Fig. 6. n.s., not significant; *, p<0.05; **, p<0.005; ***, p<0.0005.

Re-expression of wild-type SF3B1 had converse effects on splicing, either partially rescuing splicing, or for *RNF10,* increasing exon inclusion:skipping ratio beyond native levels (**Fig. 6**). Disrupting all SF3B1 ULMs with tryptophan-to-alanine mutations severely penalized the ability of the SF3B1 noULM variant to enhance the exon inclusion:skipping ratio, in agreement with its lack of binding to UHM splicing factors (23,35). Restoring only ULM5, the preferred binding site of U2AF2, partially restored splicing. The SF3B1 ULM5mut variant, in which all ULMs except ULM5 were preserved, restored splicing similarly to the WT SF3B1. This result is consistent with the ability of SF3B1 ULM5mut to bind and coimmunoprecipitate with U2AF2, yet leaves open the possibility of some ULM – UHM redundancy (e.g., a potential ability of PUF60 to substitute for U2AF2 (36)). Altogether, these results demonstrate the importance of the U2AF2 UHM and SF3B1 ULMs for pre-mRNA splicing functions in cells.

### SF3B1 and SF1 regulate splicing of U2AF2-sensitive transcripts that are mostly distinct

The U2AF2 UHM interacts with SF1 prior to SF3B1 during spliceosome assembly (6,7,9,10), raising the possibility of some redundancy in the functions of SF1 and SF3B1 for pre-mRNA splice site selection. To distinguish potential overlapping functions of the two U2AF2 partners, we compared alternative splicing of the representative, U2AF2-responsive *THYN1*, *SAT1, INTS13,* and *RNF10* transcripts following siRNA-mediated reduction of U2AF2, SF1, or SF3B1 (**Fig. 7**). In contrast with U2AF2 and SF3B1, SF1 had little or no significant effect on splicing of these transcripts, in agreement with its previously noted selective requirement for pre-mRNA splicing (8,37,38).

To more comprehensively examine an unbiased set of transcripts, we leveraged RNAseq datasets available from the ENCODE project (39–41) for U2AF2, SF3B1, and SF1 knockdown in K562 erythroid leukemia and HepG2 hepatocellular carcinoma cell lines (**Fig. 8**, **Table S4**). We identified alternative splicing events using STAR Aligner (42) and DESeq (43) with custom Python scripts as described previously (44,45) (Experimental Procedures). We then used DEXseq (65) to quantify differential splicing.

**Figure 8.**
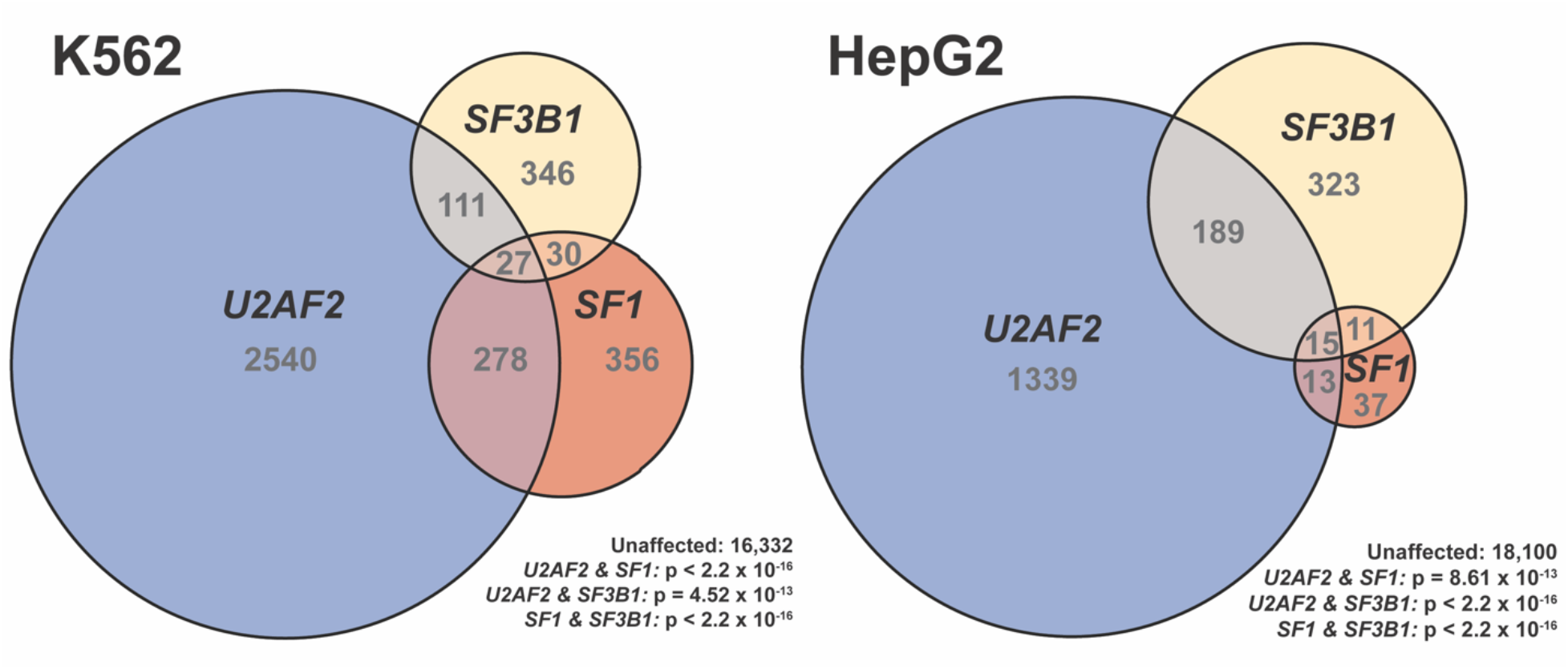
U2AF2, SF3B1, and SF1 co-regulate a subset of transcripts genome-wide. Euler diagrams showing the number of overlapping and unique alternative splicing events that exhibited statistically-significant changes upon RNAi knockdown of U2AF2, SF3B1, or SF1 relative to control shRNAs from the ENCODE project. Experiments performed in K562 erythroid leukemia cells are shown in the left diagram, and those in HepG2 hepatocellular carcinoma cells are on the right. The number of events belonging to each region is indicated. All pairwise-comparisons showed significantly greater overlap than expected by chance. For each cell line, the total number of unaffected events assayed, and the p-values for each overlap (Fisher’s Exact Test, one-sided) are shown.

More than a thousand U2AF2-sensitive splicing events were identified for each cell line. A lesser but still substantial number of differential splicing events were identified for the SF3B1-knockdown samples. In contrast, SF1-associated differential splicing events were infrequent in HepG2, consistent with a conditional, kinetic role for SF1 in metazoan BPS selection (8,37,38,46). A nearly ten-fold increase in the number of SF1-associated splicing changes in K562 cells compared to HepG2 further suggested that SF1-responsive alternative splicing is highly dependent on the cellular context. The modulated transcripts largely differed between SF3B1 compared to SF1 knockdowns for either cell line. Significant subsets of U2AF2-responsive transcripts also were regulated by either SF3B1 or SF1. With the notable exception of a subset of splicing events in K562 that are responsive to all three knockdowns, including almost 20% of SF3B1-responsive events, the SF3B1/U2AF2 or SF1/U2AF2-responsive subsets showed little overlap. These observations suggested that for the majority of splicing events, SF3B1 and SF1 contribute differently to cellular pre-mRNA splicing and the splicing-functions of U2AF2, such that some introns are more dependent on one factor or the other for efficient splicing.

## Discussion

Here, we determined the crystal structure of a cognate U2AF2 UHM – SF3B1 ULM5 complex. We show that this interface is important for U2AF2 – SF3B1 association in cell extracts and regulation of pre-mRNA splicing, whereas previous studies have been limited to the interactions of the purified protein domains.

Structurally, the U2AF2-bound SF3B1 ULM5 conformation is unique among UHM-bound SF3B1 ULM structures (**Fig. 2**). The conformational differences appear to result from an atypical lysine (K453) at the central position of the U2AF2 R**X**F motif, which differs from a bulky aromatic residue of other UHMs. An SF3B1 D339 side chain packs against the hydrophobic portion of U2AF2 K453. Accordingly, a K453A mutation significantly decreases the binding affinity of U2AF2 for the SF3B1 ULM5 (**Fig. 3**). An SF3B1 TP motif, which is a consensus site for CDK-phosphorylation, is located near the U2AF2 α-helical surface and distant from K453 in the UHM RXF motif. By contrast, the SF3B1 TP motif stacks with an aromatic central residue in the RXF motif of nearly all other UHMs. These distinct interactions raise the possibility that SF3B1 phosphorylation differently regulates its association with U2AF2 compared to other UHM-containing partners.

We established that an intact UHM is necessary for U2AF2 to associate with SF3B1 in human cell extracts (**Fig. 4**). This result is a refreshing confirmation that the long-known interaction between the purified U2AF2 UHM and the SF3B1 ULMs (23–25) is relevant for association of the fulllength factors in cells. The high affinity of the purified U2AF2 UHM for a minimal SF3B1 ULM5 (**Table S1**) further reinforced the case for a cognate interaction between these regions. Nevertheless, the binding affinity of U2AF2 for SF3B1 is moderate compared to SF1 (30). Other factors are likely to enhance and regulate U2AF2 specificity for SF3B1 *versus* SF1 in the context of the full-length, spliceosome-associated proteins, including pre-mRNA interactions by the U2 snRNA and SF3B1 HEAT repeats, dynamic RNA unwindases, and kinases/phosphatases.

Altogether, our results show that the interface residues of the U2AF2 UHM are important for splicing of a minigene prototype, and that the SF3B1 ULMs contribute to representative alternative pre-mRNA splicing events (**Fig. 5–6**). In principle, the consequences of disrupting the U2AF2 UHM and SF3B1 ULMs in these systems could result from interactions with other partners, e.g., SF1 for U2AF2 or other UHM-containing splicing factors such as Tat-SF1 for SF3B1 (22). However, we note that roles for SF1 or Tat-SF1 in pre-mRNA splicing are conditional and rare (8,38,47). Indeed, reduced SF1 levels had little effect on splicing of the U2AF2-responsive splice sites examined here (**Fig. 7**). Moreover, the graded restoration of splicing following ablation of all ULMs, only ULM5, or all but ULM5 (**Fig. 6**) agreed with the preference of U2AF2 to bind SF3B1 ULM5 while retaining the capacity to bind other ULMs (reference (23) and **Fig. 1**). Therefore, we believe that the impact of the U2AF2 UHM and SF3B1 ULMs on pre-mRNA splicing observed here is likely to arise in part or full from disrupting the U2AF2 – SF3B1 complex.

On a transcriptome-wide scale, we documented numerous, U2AF2-responsive splicing events, in agreement with a central role for U2AF2 in identifying the major class of 3’ splice sites (48), as well as previous findings of ubiquitous U2AF2 CLIP-seq sites (49,50). The lower, but still significant number of SF3B1-responsive splicing events was consistent with decreased availability of a spliceosome subunit that targets the intronic BPS (51,52), which could generally penalize splicing of sensitive introns or select an alternative BPS without detectably switching the splice site (e.g., (53)). We documented a similar number of SF1-responsive as SF3B1-responsive splicing events in K562 cells, but very few SF1-responsive splicing events in HepG2 cells. This finding reinforces conditional and cell type-dependent roles for SF1 in cellular pre-mRNA splicing (8,37,38). A substantial subset of splice sites that are responsive to both U2AF2 and SF3B1 knockdown, which are largely separate from the sites sensitive to both U2AF2 and SF1 depletion, underscores a distinct, functional relationship between U2AF2 and SF3B1.

In summary, we have demonstrated that the UHM – ULM interface is important for U2AF2 – SF3B1 association and provided evidence for its functional contribution to pre-mRNA splicing. The singular conformation of the U2AF2-bound SF3B 1 ULM5 diversifies known modes of UHM – ULM interaction, and suggests that phosphorylation of the SF3B1 TP motifs may regulate U2AF2 differently compared to other UHM-containing partners. Altogether, these results lay a groundwork for future expansions in our understanding of the structural and functional distinctions among UHM-containing proteins and their dynamic associations with the SF3B1 ULMs.

## Experimental procedures

### DNA constructs

All protein and peptide sequences correspond to the human homologues. The U2AF2 UHM construct was described previously (20,23). The U2AF2^12UL^ construct includes residues 141 to the C terminus (residue 471) of NCBI RefSeq NP_009210. The SFB1^147-462^ construct includes residues 147 – 462 of NCBI RefSeq NP_036565. The full-length SF3B6 construct matches to NCBI RefSeq NP_057131. For transfections, the plasmids encoding full-length U2AF2 and SF3B1 were described previously (32,54,55). Structure-guided mutations were introduced by Genscript.

### Preparation of purified proteins

All proteins were expressed using the pGEX-6p vector, purified by glutathione affinity, proteolytic cleavage to remove the GST-tag, ion exchange, and a final size exclusion chromatography step in 50 mM NaCl, 25 mM HEPES pH 7.4, 0.2 mM TCEP through a Superdex-75 column (Cytiva). Synthetic peptides were purchased with >98% purity (Biomatik Corp.).

### Isothermal titration calorimetry

A MicroCal VP-ITC (Malvern Panalytical) was used to inject 28 aliquots of 10 μl each at a rate of 2 sec μl^-1^ separated by a 4 min relaxation time into the sample cell. The experiments were run at 30 °C, 15 μcal s^-1^ reference power, and with constant stirring at 307 rpm. Concentrations were typically 5 μM SF3B1 in the sample cell and 50 – 100 μM U2AF2 in the syringe. The isotherms were corrected for the heats of dilution by subtracting the last three data-points of the saturated region, then fit using Origin^R^ v7.0 (Malvern). The ITC results are detailed in **Table S1** and isotherms are shown in **Fig. S1**.

### Crystallization and structure determination

The U2AF2 UHM – SF3B1 ULM5 complex was crystallized by the hanging drop vapor diffusion method at 4 °C from 20.5 mg/ml U2AF2 UHM in the presence of a 1.5-fold molar excess of SF3b155 ULM5 peptide. The reservoir solution contained 1 M sodium citrate tribasic dihydrate pH 5.6, 15% v/v 2-propanol, 20% v/v PEG 4000. For cryo-protection, crystals were sequentially transferred to reservoir solution supplemented with 10% v/v glycerol, then flash-cooled in liquid nitrogen. Crystallographic data sets at 100 K were collected by remote data collection at the Stanford Synchrotron Radiation Light (SSRL) source Beamline 12-2 (56). The data was processed using the SSRL AUTOXDS script (A. Gonzalez and Y. Tsai), which implements XDS (57) and CCP4 packages (58). The structure was determined by molecular replacement using Phaser (59) with the U2AF2 UHM from the SF1 complex (PDB ID: 4FXW) as the search model (20). A top solution with LLG of 1253 and a top TFZ-score of 26.7 showed clear electron density for the SF3B1 peptide bound to both copies of the U2AF2 UHM in the crystallographic asymmetric unit (feature-enhanced electron density maps (60) shown in **Figure S2**). The structure was refined in Phenix.refine (61) and manually adjusted in Coot (62). The crystallographic data collection and refinement statistics are reported in **Table S2**.

### Cell culture and transfections

Human embryonic kidney epithelial cells (HEK 293T, ATCC^®^ CRL-3216^TM^) were maintained at 37 °C in a humidified atmosphere containing 5% CO2 as described previously (32). Cells were transfected in six-well plates at 50-60% confluency with the indicated siRNAs and/or DNA plasmids, using jetPRIME^R^ (Polyplus-transfection SA) as instructed by the manufacturer. For experiments with the *pyPY* minigene, a stable 293T cell line expressing the *pyPY* transcript was transfected with plasmids expressing U2AF2 variants, then cells were harvested 24 h after transfection as described (32). For knockdown experiments, cells were transfected with 25 nM of Stealth^TM^ siRNAs (Thermo Fisher Sci.), targeting either U2AF2 (catalog nos. HSS117616, HSS117617), SF1 (catalog nos. HSS187735, HSS144483), SF3B1 (catalog nos. HSS146413, HSS146415) or a “Lo GC” control (catalog no. 12935200) and harvested two days after transfection. For “rescue” experiments with the SF3B1 variants, the samples were harvested two days after co-transfection of the siRNAs and plasmid DNAs.

### Co-immunoprecipitation

HEK 293T cells were transfected in 10 cm plates with combinations of wild-type or mutated plasmids encoding HA-tagged U2AF2, FLAG-tagged SF3B1, or empty vector control (pCMV5-XL6). Since the ULM mutations appeared to alter ^FLAG^SF3B1 expression, the amounts of transfected SF3B1 constructs and empty control vector were adjusted to achieve similar levels of ^FLAG^SF3B1 variants among the co-immunoprecipitation inputs, while maintaining equivalent amounts of total transfected DNA. After 24 h, cells were harvested and lysed in IP buffer (50 mM Tris pH 8.0, 75 mM NaCl, 5% v/v glycerol, 10 mM CaCl2, cOmplete^TM^ EDTA-free protease inhibitor (Sigma-Aldrich), β-glycerophosphate, 0.5 mM DTT) plus 0.5% v/v Triton X-100. Resuspended cells were sheared then centrifuged to remove debris. Equal amounts of total protein (DC Protein Assay, Bio-Rad) were used for the immunoprecipitation reactions. First, a fraction of each sample was set aside as an input control. Then, 1 mg of each of the remaining lysates was diluted four-fold with IP buffer plus 0.1% v/v Triton X-100 and incubated for 2 hr at 4 °C with protein G-Sepharose (GE Healthcare catalog no. 17061801), pre-bound to HA-specific antibody (rabbit anti-HA from Sigma-Aldrich, catalog no. H6908) (3.5 μg antibody per reaction). Beads were collected by centrifugation and washed six times with IP buffer before analysis by SDS-PAGE.

### Immunoblotting

For total protein analysis, harvested cells were lysed in 50 mM Tris pH 8.0, 10 mM EDTA, 1% w/v SDS, 1 mM DTT, phosphatase inhibitors, and protease inhibitors. Total protein concentrations were measured (DC Protein Assay, Bio-Rad) and equal amounts of protein were loaded per lane of SDS-PAGE. Separated proteins were transferred to polyvinylidene difluoride (PVDF) membranes and immunoblotted with antibodies specific for SF3B1 (Abcam, catalog no. ab170854), SF1 (Bethyl Laboratories, catalog no. A303-213A), U2AF2 (Sigma-Aldrich, catalog no. U4758), FLAG (Sigma-Aldrich, catalog no. F1804), HA (Sigma-Aldrich, catalog no. H6908), or GAPDH (Cell Signaling, catalog no. 14C10), all diluted 1:1000 v/v with 5% w/v dry milk in TBS-T. Secondary antibodies included anti-rabbit IgG horseradish peroxidase (Invitrogen, catalog no. 31460) or antimouse horseradish peroxidase (Invitrogen, catalog no. 31340). The chemiluminescence signal from Clarity™ Western ECL substrate (Bio-Rad, catalog no. 170-5061) was detected on a Chemidoc™ Touch Imaging System (Bio-Rad).

### RT-PCR of minigene and endogenous gene transcripts

The protocols used for RT-PCR were described previously (32). Briefly, total RNA was isolated from harvested cells and DNase I-treated using the RNeasy kit (Qiagen). The cDNAs were synthesized using random primers and Moloney murine leukemia virus RT (Invitrogen). The RT-PCR products were separated on a 2% w/v agarose-TBE gel, stained with ethidium bromide, and visualized using a Gel Doc XR+ gel documentation system (Bio-Rad). The band intensities of three technical replicates were quantified and background-corrected using ImageJ {Abramoff, 2004 #2213}, and are representative of multiple biological replicates. The primer sequences are listed in **Table S3**.

### Bioinformatics and statistical analysis

#### RNAseq read alignment

RNAseq analysis was performed as previously described (44,45). FASTQ files were downloaded from the ENCODE project site (https://www.encodeproject.org/; sample accession numbers are listed in **Table S4**). These included eight control replicates treated with the control shRNA and two biological replicates for each knockdown (SF3B1, U2AF2, SF1) from both K562 erythroid leukemia and HepG2 hepatocellular carcinoma cell lines. Reads were mapped to the Hg38 human reference genome using STAR Aligner (42). Mapping statistics are listed in **Table S4**.

#### Identification of alternative splicing events

Splice junctions determined by STAR mapping were combined for all samples and collapsed into a non-redundant set of introns. Alternative and constitutive intron classifications were performed using custom Python scripts, and are agnostic with regard to existing annotations other than known gene boundaries (44,45). The workflow takes a set of intron coordinates, assigns them to a gene, and divides them into subgroups based on overlapping coordinates. If no overlapping introns exist for a given intron, it is assigned to the *constitutive* class. The subgroups containing overlapping introns are assigned a splicing classification if the start and end coordinates of all of the constituent introns fall into a pattern representing a known splice type *(cassette, mutually exclusive, alternative 5’ splice site, alternative 3’ splice site).* For identification of detained introns, STAR-mapped reads were filtered to remove intron-spanning reads and reads aligning to annotated exons or repeat RNAs. To remove polyadenylation sites within introns that might contribute to false-positive DI identification, the genome-wide coordinates of known polyadenylation sites were extracted from the GENCODE annotation (63), and introns containing these sites were not considered in assignment of DI status. Finally, alternative splicing annotations from the steps above were used to further filter out any introns that might contain exons or other introns to produce a non-overlapping set of introns spanning each gene locus. Mapped and filtered reads were assigned to introns using Bedtools (64). For each gene, the sum of normalized intronic read counts was used to allocate reads to individual introns under a null model based on their length and mappability. A variance stabilizing transform based on the square root of intron effective length adjusted by RNAseqread length (√(Ld)where L=mappability adjusted intron length, d = RNAseq read length) was then used to weight individual introns. The sum of normalized intronic reads per gene in each RNAseq replicate was then partitioned and allocated to each intron proportional to its weight. This results in an *in silico* null model replicate corresponding to each RNAseq replicate. Differential analysis using DESeq (43) was then used to determine introns enriched in read coverage (in the RNAseq replicates) compared to the *in silico* null model replicates using an FDR-adjusted p value threshold of 0.01 and fold change threshold of 2.

#### Differential splicing analysis

Annotated alternative and constitutive exons were used as an input to generate an ‘exon part’ gtf that was compatible with DEXSeq, using the script dexseq_prepare_annotation.py (65). Reads were counted from mapped bam files using the counting script dexseq_count.py to generate count tables for each exon part. Differential expression of the alternative splicing events and detained introns was then determined using standard DEXSeq analysis with a padj. <0.05 as the cutoff for significant changes, comparing the eight control replicates against each of the two replicate sets for each knockdown. Overlapping events that were differentially spliced in each of the knockdowns were then counted, and the proportional Euler diagrams were produced in R using the package eulerr (66). Statistical significance of overlaps was determined using one-sided Fisher’s Exact Test calculated in R version 3.6.3 (67).

## Supporting information

This article includes Tables S1-S4 and Figs. S1-S4.

## Data availability

Atomic coordinates and structure factors of U2AF2 UHM bound to SF3B1 ULM5 (accession code 7SN6) have been deposited at the Protein Data Bank (http://wwpdb.org).

## Acknowledgments

We are grateful to Maria Carmo-Fonseca (U. Lisbon, Portugal) for providing the *pyPY* and wild-type HA-U2AF2 plasmids, Esther Obeng and Benjamin Ebert (Dana-Farber) for the SF3B1 template plasmid, and Steven Horner and Justin Leach for contributions to ITC. Data from the ENCODE consortium were courtesy of Brenton R. Graveley (U. Conn. Health).

## Abbreviations

The abbreviations used are:

BPS: branch point sequence;
CDK: cyclin-dependent kinase;
cryoEM: cryo-electron microscopy;
ENCODE: encyclopedia of DNA elements;
HEK: human embryonic kidney;
IP: immunoprecipitation;
PDB: protein data bank;
Py: polypyrimidine;
RMSD: root mean square deviation;
RRM: RNA recognition motif;
snRNP: small nuclear ribonucleoprotein;
UHM: U2AF homology motif;
U2AF: ligand motif.

