## Supporting information for "A UHM – ULM interface contributes to U2AF2 and SF3B1 association for pre-mRNA splicing"

Contents:

Table S1: Isothermal titration calorimetry results
Table S2: Crystallographic data and statistics
Table S3: Primer sequences
(Table S4 is provided in a separate Excel file: Summary of ENCODE datasets)

Figure S1: Representative isotherms for Figures 1 and 3
Figure S2: Electron density maps for SF3B1 ligands in Figure 2
Figure S3: Immunoblot for minigene transcript splicing in Figure 5
Figure S4: Immunoblots for endogenous transcript splicing in Figures 6 and 7

**Table S1.** Isothermal titration calorimetry of U2AF2 regions titrated into SF3B1 regions.

| **Interaction:** [a] | **n** | **$K_D$** (nM) | **ΔG** [b] (kcal mol$^{-1}$) | **ΔH** (kcal mol$^{-1}$) | **-TΔS** [c] (kcal mol$^{-1}$) |
|---|---|---|---|---|---|
| **Wild-type U2AF2 UHM titrated into SF3B1 ULM5 residue ranges:** | | | | | |
| 329 – 344 [d] | 0.9 ± 0.1 | 449 ± 75 | -8.8 ± 0.1 | -15.0 ± 1.4 | 6.2 ± 1.4 |
| 329 – 351 | 0.9 ± 0.1 | 108 ± 13 | -9.7 ± 0.1 | -14.0 ± 0.7 | 4.3 ± 0.7 |
| 333 – 351 | 1.0 ± 0.1 | 49 ± 2 | -10.1 ± 0.0 | -19.8 ± 0.6 | 9.6 ± 0.6 |
| 190 – 344 [d] | 2.9 ± 0.2 | 2,833 ± 23 | -7.7 ± 0.01 | -9.4 ± 0.7 | 1.7 ± 0.7 |
| **Wild-type U2AF2$^{12UL}$ titrated into SF3B1$^{147\text{-}462}$ +/- SF3B6:** | | | | | |
| – | 2.0 ± 0.1 | 797 ± 80 | -8.5 ± 0.1 | -9.4 ± 0.2 | 0.9 ± 0.3 |
| + SF3B6 | 1.8 ± 0.2 | 940 ± 120 | -8.4 ± 0.1 | -10.4 ± 1.0 | 2.0 ± 1.0 |
| **Wild-type U2AF2 UHM titrated into mutant SF3B1 ULM5 residues 333 - 351:** [e] | | | | | |
| T341A/M346A | 0.8 ± 0.1 | 38 ± 4 | -10.3 ± 0.1 | -20.1 ± 0.0 | 9.8 ± 0.0 |
| P342G | 0.8 ± 0.1 | 63 ± 12 | -10.0 ± 0.1 | -18.3 ± 0.2 | 8.3 ± 0.1 |
| **Mutant U2AF2 UHM titrated into SF3B1 ULM5 residues 333 - 351:** | | | | | |
| K453A | 0.8 ± 0.1 | 210 ± 26 | -9.3 ± 0.1 | -17.3 ± 0.1 | 8.1 ± 0.2 |
| E394K/E397K [f] | *No heats of binding detected* | | | | |
| **Wild-type U2AF2 UHM titrated into SF1 residue ranges:** | | | | | |
| 1 – 255 [d] | 0.9 ± 0.1 | 12 ± 5 | -11.0 ± 0.3 | -17.5 ± 0.7 | 6.5 ± 0.4 |
| 13 – 28 (ULM) [d] | 0.9 ± 0.1 | 24 ± 4 | -10.6 ± 0.1 | -11.1 ± 0.4 | 0.5 ± 0.5 |

[a] Average values and standard deviations of three independent titrations. n, apparent stoichiometries. U2AF2$^{12UL}$ is residues 141 – 471 (C-terminus) of isoform b and includes an N-terminal α-helix of RRM1, RRM1, RRM2, and the UHM; U2AF2 UHM is residues 361-461 of isoform b and is identical to residues 365-475 of isoform a; SFB1$^{147\text{-}462}$ is residues 147-462 and includes the ULMs and SF3B6-binding site.

[b] Calculated using the equation $\Delta G = -RT \ln (K_A)$ where T=303 K.

[c] Calculated using the equation $-T\Delta S = \Delta G - \Delta H$.

[d] Reproduced from Thickman *et al* (2006) *J. Mol. Biol.* v356:664. Experiments were conducted with the same protein preparation methods, calorimeter model, and buffer compositions (within measurement error).

[e] Please refer to 333-351 as the matching WT counterpart for comparison with SF3B1 ULM5 mutants.

[f] No binding was detected during titration of 50 μM SF3B1 ULM5 (residues 333-351) into 5 μM U2AF2 UHM E394K/E397K mutant.

### Table S2. X-ray data collection and refinement statistics.

Values in parentheses refer to the highest resolution shell (1.84 - 1.80 Å). For $R_{free}$, 9% of the data (2,008 reflections) were randomly-selected for exclusion from all stages of refinement.

$R_{merge} = \sum_{hkl}\sum_i|I_i - \langle I\rangle|/\sum_{hkl}\sum_i |I_i|$ , where $I_i$ is an intensity I for the ith measurement of a reflection with indices hkl and <I> is the weighted mean of all measurements of I.

$R_{p.i.m.} = \sum_{hkl} (1/(n-1))\sum_i|I_i - \langle I\rangle|/\sum_{hkl}\sum_i |I_i|$ where n is the number of observations of the intensity $I_i$.

$CC_{1/2}$, correlation coefficient between intensities of random half-dataset.

$R_{work} = \Sigma_{hkl}||F_{obs}(hkl)|-|F_{calc}(hkl)||)/\Sigma_{hkl}|F_{obs}(hkl)|$ for the working set of reflections.

$R_{free}$ is $R_{work}$ for ~7% of the reflections excluded from the refinement. No cutoff was applied in the refinement.

The MolProbity score was calculated using the program MolProbity.

| ULM5-bound U2AF2 UHM | |
|---|---|
| **Data collection** | |
| Wavelength (Å) | 1.03 |
| Resolution range (Å) | 37.97 - 1.80 |
| Space group | *C* 2 |
| Unit cell lengths (Å) | 76.2, 63.2, 49.4 |
| Unit cell angles (°) | β = 94.4 |
| Total reflections | 77069 |
| Unique reflections | 21337 |
| Multiplicity | 3.6 (3.6) |
| Completeness (%) | 98.4 (96.8) |
| Mean I/σ(I) | 16.7 (3.0) |
| $R_{merge}$ (%) | 3.8 (49.9) |
| $R_{p.i.m.}$ (%) | 2.3 (30.4) |
| $CC_{1/2}$ (%) | 99.0 (92.0) |
| **Refinement** | |
| $R_{work}/R_{free}$ (%) | 18.0 / 21.7 |
| U2AF2 UHM in ASU | 2 |
| SF3B1 ULM5 in ASU | 2 |
| No. non-hydrogen atoms: | |
| U2AF2 UHM | 1656 |
| SF3b1 ULM5 | 230 |
| 2-Propanol | 4 |
| Waters | 71 |
| R.m.s. Bonds (Å) | 0.01 |
| R.m.s. Angles (°) | 1.0 |
| Ramachandran (%) favored/allowed/outlier | 99.5/0.5/0 |
| Clashscore (percentile) | 3.7 (98) |
| MolProbity score | 1.16 |
| <B-factor> (Å$^2$): | 47.5 |
| U2AF2 UHM | 46.2 |
| SF3B1 ULM5 | 56.8 |
| 2-Propanol | 76.0 |
| Water | 46.9 |
| **PDB Code** | 7SN6 |

**Table S3. Sequences of primers used for RT-PCR (5´-to-3´)**

| Gene Name | Forward Primer | Reverse Primer |
|---|---|---|
| *pyPY* | TGAGGGGAGGTGAATGAGGAG | TCCACTGGAAAGACCGCGAAG |
| *THYN1* | CCCCATTATGACCCATCTAGCA | GCCCATTCCAGCAGCAGTAT |
| *SAT1* | TTTGGAGAGCACCCCTTTTA | CTCATCACGAAGAAGTCCTCAA |
| *RNF10* | GGCCCTCTCCATTTCTCCTC | TTATCACTCTCCCCGTCGCT |
| *ASUN* | GTGAAGGATGTGGAGGAAGAGTT | TCACAATAACACTGGCTAATGGA |

**Table S4. Summary of ENCODE datasets.**

Please see separate Excel file.

**Figure S1.** Representative isotherms of U2AF2 regions titrated into SF3B1 regions. Data points (squares) were fit with identical-site binding models (solid lines). *A – B,* wild-type U2AF2 UHM and SF3B1 ULM5 peptides of the given residue ranges; *C – D,* nearly fully length U2AF2 and SF3B1, either alone or co-purified with SF3B6; *E – F*, U2AF2 UHM mutants and SF3B1 ULM5, residues 333-351; *G – H*, wild-type U2AF2 UHM and mutants of SF3B1 ULM5, residues 333-351. The apparent dissociation constants are inset (average values and standard deviations of three independent titrations).

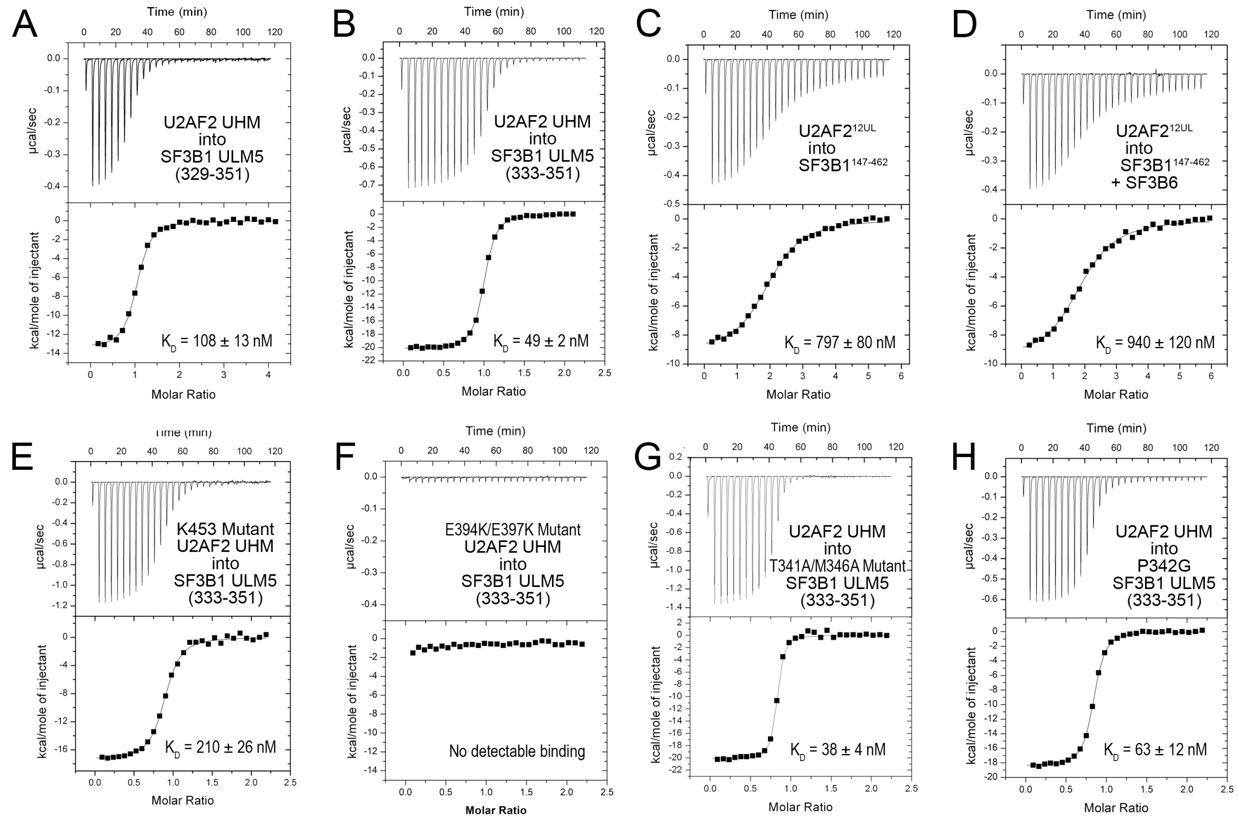

**Figure S2.** Crystal structure of U2AF2 UHM – SF3B1 ULM5. *A,* Crystal packing showing contacts between the U2AF2 UHM (dark blue) ordering the C-terminus of the symmetry-related ULM5 in Complex B (orange, ULM5´). The UHM/ULM of complex A are colored light blue/yellow. The asymmetric unit is enclosed in a dashed rectangle and the labels of the symmetry-related complex are primed. *B*, Reduced-bias electron density maps contoured at 1 σ confirm SF3B1 ULM5 is bound to both copies of the U2AF2 UHM in the crystallographic asymmetric unit.

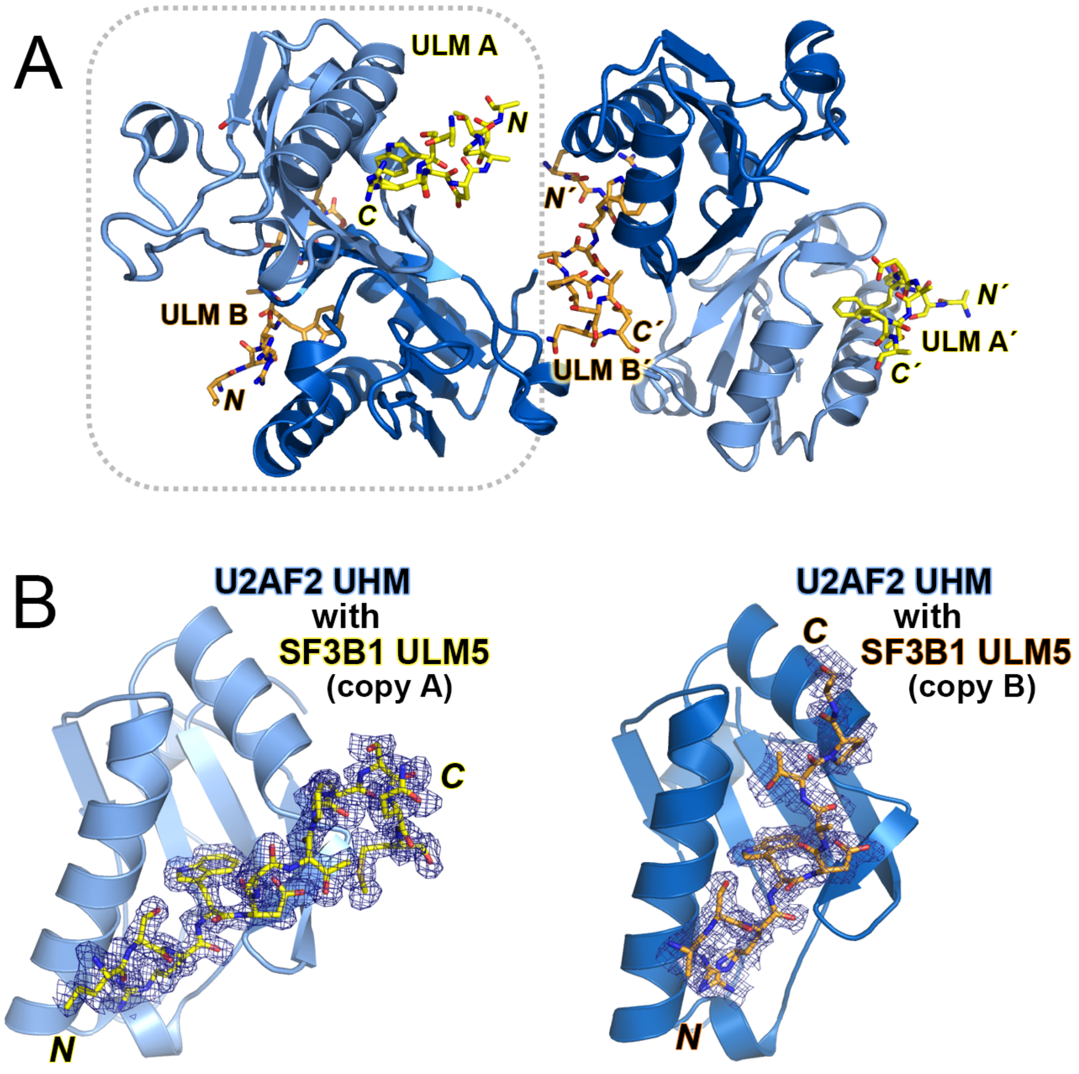

**Figure S3.** Immunoblots of HEK 293T expressing the *pyPY* minigene and transfected with wild-type (WT) or mutant (E394K/E397K) U2AF2 (65 kDa) for Figure 5. The lower portion of the blot was probed for GAPDH (37 kDa) as a loading control. Colorimetric images of the molecular size standards (STD) are overlaid on the chemiluminescent blots.

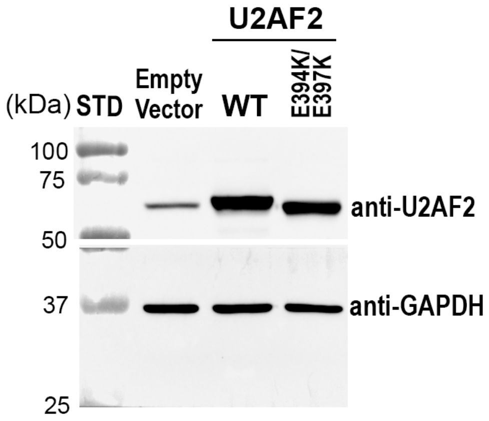

**Figure S4.** Immunoblots of the HEK 293T samples transfected with *A,* SF3B1 siRNA #2 (siSF3B1 #2) and SF3B1 variants for the RT-PCR experiments in Figure 6 and *B,* siRNAs targeting SF3B1 (145 kDa), SF1 (69 kDa; the second, supershifted band may be post-translationally modified or an isoform), or a loGC control siRNA for the RT-PCR experiments in Figure 7. Intensities of saturated bands (red) are above the linear range of detection. Colorimetric images of the molecular size standards (STD) are overlaid on the chemiluminescent blots.

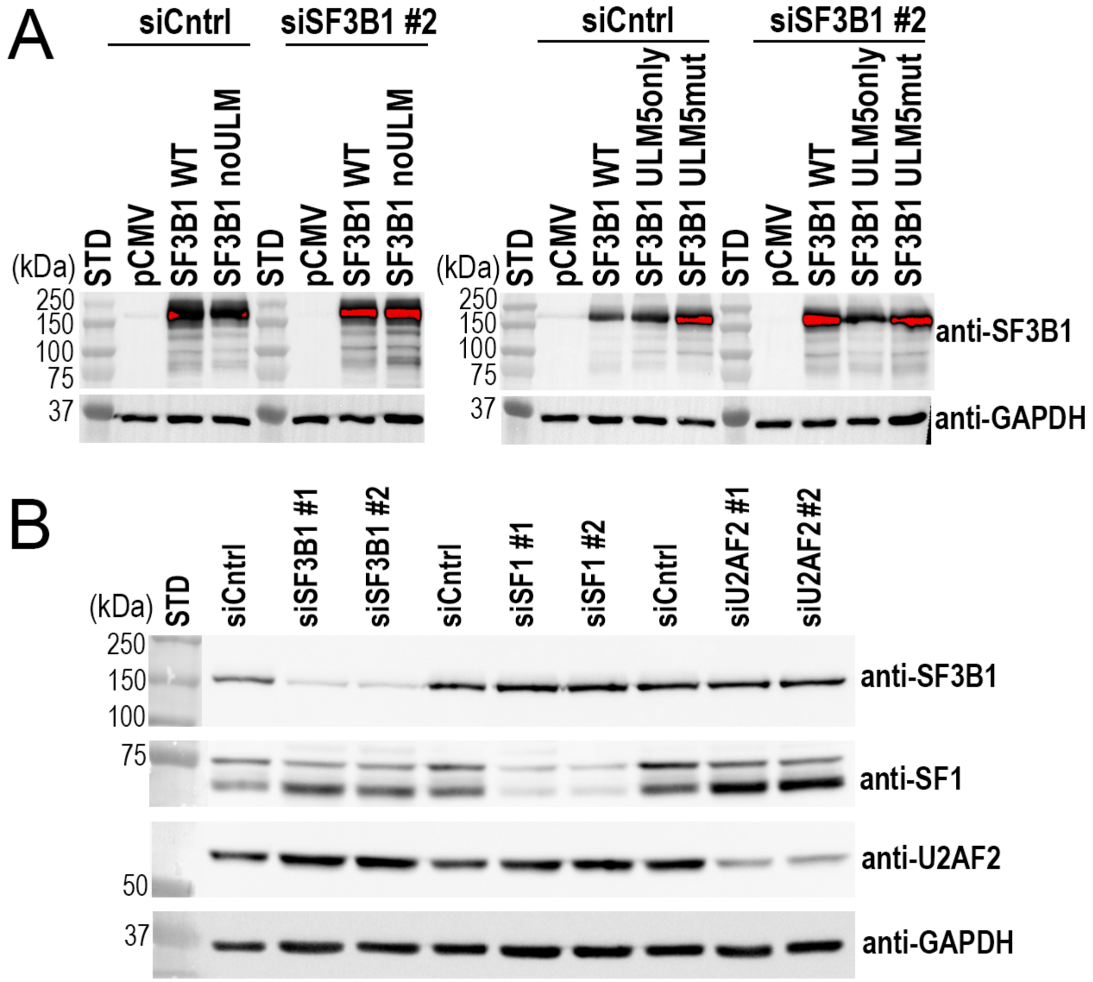

### Supplementary References
